# Development of the 4TREE SNP array, a forest multispecies array to enhance European Breeding and conservation programs in pine, poplar and ash

**DOI:** 10.64898/2026.03.21.711309

**Authors:** R. Guilbaud, F. Bagnoli, S. Ben-Sadoun, C. Biselli, C. Buret, J. Buiteveld, L. Cattivelli, P. Copini, J. Drouaud, D. Esselink, A. Fricano, V. Benoit, L.J. Kelly, L. Kodde, C.L. Metheringham, S. Pinosio, O. Rogier, V. Segura, I. Spanu, G. Tumino, R.J.A. Buggs, S. C. González-Martínez, L. Vietto, G. Nervo, V. Jorge, A. Dowkiw, M.J.M. Smulders, L. Sanchez, G.G. Vendramin, C. Bastien, P. Faivre Rampant

## Abstract

Within the framework of the European Adaptive BREEDING for Better FORESTs project (B4EST, https://b4est.eu/), we have developed genotyping tools for Poplar, Ash, and Pine forest tree species. SNP arrays are attractive genotyping tools because of the user-friendly genotype calling system and the robust transferability among laboratories.

Here we describe the development of an Axiom SNP array for *Pinus pinaster* (13,407 SNPs), *Pinus pinea* (5,671 SNPs), *Poplar* spp. (13,408 SNPs), and *Fraxinus* spp. (13,407 SNPs) based on a two-step process. We first assembled a high-density (>100,000 SNPs/species) screening array that served to test a large panel of candidate SNPs on a diversity panel involving at least 120 individual trees per species or species group. In the second step, we selected and combined the most informative SNPs to build the final 50,000 SNP 4TREE array.

This approach resulted in high genotyping success rates, including for species lacking previously validated high-quality SNP resources. The 4TREE SNP array provides a valuable and transferable genomic tool to support genomic prediction, breeding, and adaptive management of forest tree species.

## INTRODUCTION

Forest trees are perennial species characterized by long juvenile phases, late flowering, and large body size at maturity. When such species are the focus of genetic improvement programs, these biological characteristics have a negative impact on the breeding cycle, making them costly in terms of time, space, and resources. These constraints limit the capacity of conventional breeding programs to rapidly respond to emerging challenges such as climate change, novel pests, and diseases. In this context, Genomic Selection (GS) has emerged as a powerful approach to accelerate genetic gain by predicting phenotypic performance from genome-wide marker information, even with limited phenotyping efforts. It has proven effective in livestock and crops and is starting to evolve from proof of concept to reality in trees (Lebedev et al. 2020; Pegard et al. 2020; Grattapaglia 2022a; Gamal-El-Dien, 2024; Sharma et al. 2025). This approach requires genotyping tools with a high number of available molecular markers spanning the whole genome. Such tools have been created since the early 2000s in the wake of the development of high-throughput genomics. There are currently two main categories of single nucleotide polymorphism (SNP)-based genotyping platforms: classical SNP arrays and Genotyping by Sequencing (GBS). Several possible approaches are available in this latter category, including Whole Genome Sequencing (WGS), reduced representation sequencing (Elshire et al., 2011; Peterson et al, 2012), and targeted sequencing (Tripodi, 2023). While reduced representation sequencing methods offer cost-effective solutions, they frequently suffer from inconsistent variable marker representation, high levels of missing data, and limited inter-laboratory reproducibility (Scheben et al. 2017; Rasheed et al. 2017; Torkamaneh et al. 2018; Scaglione et al. 2019). Although WGS and targeted sequencing approaches provide more comprehensive and consistent coverage with lower missing data rates, their higher costs can limit their application in large-scale breeding programs, especially for large genomes like those of conifers (Pavan et al. 2020; Candotii et al. 2023; Aguire et al. 2024). In contrast, SNP arrays (or so-called DNA chips) combine cost-effectiveness with reliability, making them attractive genotyping tools, notably because of their user-friendly genotype reference free calling system, standardized marker sets, high call rates, robust transferability between laboratories, and straightforward data integration over time. Two commercial alternative array techniques are available: Infinium/Illumina® and Affymetrix/Thermo Fisher Axiom® genotyping arrays. These two designs use different technologies and chemistries, yet the SNP call rate concordance between individuals tested by both technologies may be as high as 97% (Howard et al. 2021). Both technologies have proven successful for genetic diversity studies, linkage mapping, Genome-Wide Association Studies (GWAS), and GS in several forest tree species (Table S1). Despite these advantages, the availability of validated SNP array resources remains limited for many economically and ecologically important forest trees.

The European Adaptive BREEDING for Better FORESTs project (B4EST, https://b4est.eu/) aimed at developing adaptive breeding strategies for productive, sustainable, and resilient forests under climate change, with a focus on the most economically, ecologically, and socially important tree species in Europe. One objective is to marshal expertise for genomic prediction in key European forest trees such as pine, spruce, poplar, and ash. To this aim, in the framework of this project, three SNP arrays were developed: two 50k-SNP-single-species arrays, one for *Pinus sylvestris* (Kastally et al. 2022) and one for *Picea abies* (Bernhardsson et al. 2021), and a multi-species array for *Pinus pinea*, *Pinus pinaster*, *Populus* sp., and *Fraxinus* sp. The design and evaluation of this third array is presented here.

Among the species included in the multi-species array, poplar (*Populus nigra*) and maritime pine (*Pinus pinaster*) have benefited the most from advanced genomics and genotyping tools. This has led to the first proof-of-concept genomic assessments in the context of European tree breeding programs, e.g. in *P. nigra* (Allwright et al. 2016; Faivre-Rampant et al. 2016; Scaglione et al. 2019; Pegard et al. 2020) and *P. pinaster* (Plomion et al. 2016). Poplar was the first tree genus to have its whole genome sequence made publicly available (Tuskan et al. 2006). Moreover, considerable genomic resources exist for *Populus* spp. (Porth et al. 2024). This genus is the focus of active breeding in France and Italy, especially involving *P. nigra* x *P. deltoides* hybrids (Biselli et al. 2022). *Pinus pinaster* is another species with already important contributions in assessing genomic selection (Bartholomé et al. 2016; Isik et al. 2016).

Conversely, stone pine (*P. pinea*) and ash (*Fraxinus* spp.) both face the challenge of reduced genetic diversity for a short term breeding program. In pine, this reduction is due to historical and evolutionary events (Alonso et al. 2007; Vendramin et al. 2008; Mutke et al. 2011; Jaramillo-Correa et al. 2020), whereas in ash, it is the result of recent disease outbreaks devastating European ash populations (Downie et al. 2017; George et al. 2024). Despite the urgency, these two species have not yet benefited from the development and implementation of genotyping tools.

Thermo Fisher Scientific (TF) (Thermo Fisher Scientific, Inc, Santa Clara, CA, USA) suggested a two-step process to create a robust, high quality final genotyping tool, even for difficult taxa. The first step involves a high-density (>100,000) SNP array (or so-called screening array) that serves as a benchmark for testing a large panel of candidate SNPs against a diversity panel of about 120 individuals/species. In the second step, the high-quality SNPs resulting from the screening array are recovered to build the high-quality final 50,000 SNP multispecies 4TREE array. The new multispecies SNP array developed by the B4EST consortium is the first to include SNPs specific to *P. pinea*, *P. deltoides*, and *Fraxinus* sp. (Table S1). By pooling multispecies into a single genotyping array, both SNP and samples are maximized, leading to reduced cost per species with limited research funding.

## MATERIAL & METHODS

### Screening array development

SNP discovery and selection for the screening array

### Populus spp

*P. nigra* SNP data were obtained from existing genomic resources, including WGS and RNA sequencing (RNA-Seq) datasets (Faivre-Rampant et al. 2016; Rogier et al. 2018; Pégard et al. 2019; Rogier et al. 2023). The SNP search was performed according to the procedure described in Rogier et al. (2018, 2023), resulting in a total of 26,759,614 SNPs. In addition, 8,500 high quality SNPs were retrieved from the existing Infinium array (Faivre-Rampant et al. 2016) (Figure 1).

**Figure 1:**
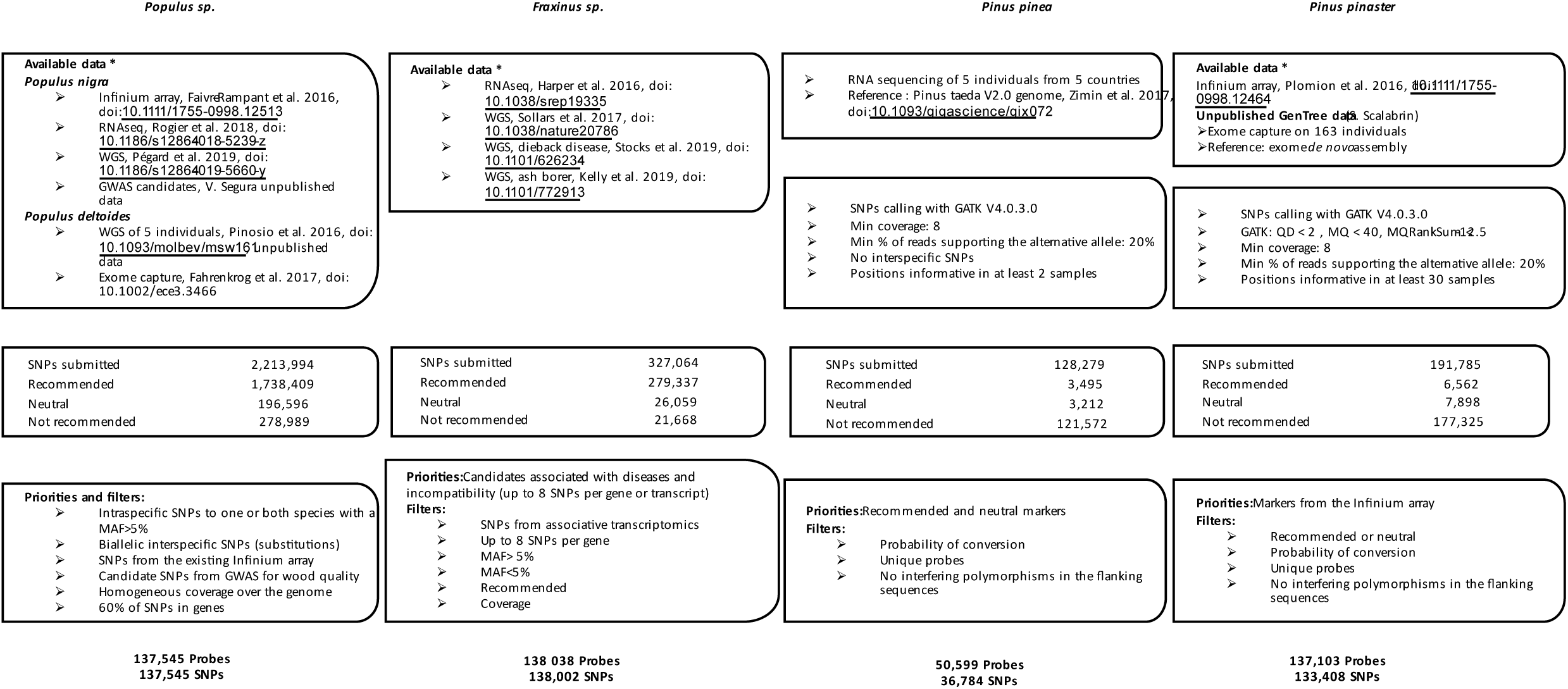
Workflow of the SNPs selection process for the screening array.

For *P. deltoides*, available genomic resources including WGS data from 5 individuals (Accession number: SRS 1 320 691, SRS 1 328 499, INRAE unpublished data) and exome capture sequencing data (Fahrenkrog et al. 2017) were used for SNP detection. The SNP calling was carried out using the same method for SNP detection in *P. nigra* (Rogier et al. 2018, 2023). Combining the different *P. deltoides* genomic datasets resulted in 887,974 SNPs (Figure 1).

The whole set of *Populus* SNPs was filtered to remove duplicates and SNPs with mismatches in their 35 bp flanking sequences. Four SNPs categories were then selected following the criteria: 1) biallelic SNPs specific to each species; 2) biallelic SNPs with alleles shared between both species; 3) substitutions, loci fixed within species but with different alleles between species; and 4) SNPs well distributed across the genome. Ultimately, a total of 2,213,994 filtered SNPs were sent to TF for quality scoring. For each SNPs, both forward and reverse probes were given p-convert values to predict the probability of SNPs conversion on the array (https://downloads.thermofisher.com/Axiom_Analysis_Suite_v_4.0.1_User_Guide.pdf). A SNPs marker/strand is recommended if: p-convert >0.6, no wobbles, and poly count= 0. In other words, a SNPs marker/strand was not recommended if it had duplicate count >0 or poly count >0 or p-convert =3. Therefore, if a marker has the same recommendation for each strand, then it was tiled on the strand with the highest p-convert value. SNPs probes with a p-convert value > 0.6 were expected to convert on the SNPs array with high probability. Among the filtered SNPs, 1,738,409 (78,5%) were successfully tiled (Figure 1, Table S2).

A selection of 137,545 probes for 137,545 SNPs markers was made for the screening array. The priority was to include intraspecific SNPs to one or both species with a minor allele frequency (MAF) > 0.05, substitutions were also included, as well as SNPs from the existing Infinium array (Faivre-Rampant et al. 2016) and candidate SNPs obtained through a GWAS (Segura, pers. com. Durufle et al. 2025). The selected SNPs were well distributed across the genome (Figure 2). Finally, the screening array combined 57,969 *P. deltoides*-specific SNPs, 66,625 *P. nigra*-specific SNPs, 12,191 SNPs that were shared by both species, and 760 substitutions. Genome assembly versions V2 and V3 were successively used as reference sequences for SNPs identification and selection. In 2022, an improved assembly (version 4.0) was released (https://phytozome-next.jgi.doe.gov/info/Ptrichocarpa_v4_1); therefore, SNPs positions were updated onto the latest assembly. Flanking sequences of selected SNPs were aligned with the BLASTn tool using the following parameters; alignment length >0,9 and % of identical match >0,95 (Figure 2).

**Figure 2:**
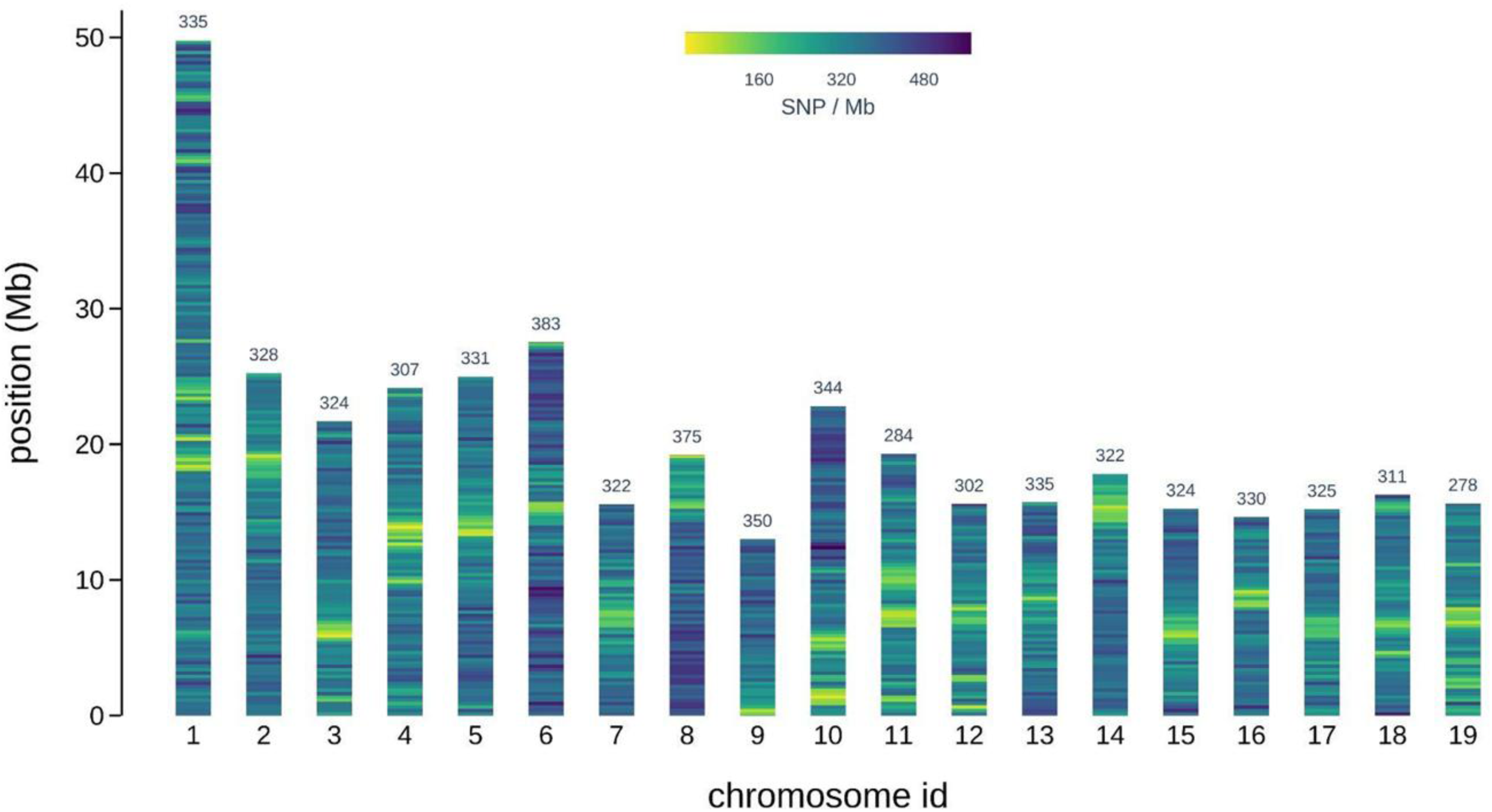
*Populus* SNPs density from the screening array on the 19 chromosomes of *Populus trichocarpa* V4.0 genome. We selected *P. trichocarpa* V4.0 for SNPs position, due to its use as a reference by the scientific community. The y-axis represents the physical distance along each chromosome, split in 250 kb windows. The different colors depict SNPs density. Over 93 % of the probes were successfully mapped along the *P. trichocarpa* v4.0 genome assembly with the BLASTn tool using the following parameters: alignment coverage >0,9 and identity >0,95. The numbers above the chromosomes are the average number of SNPs per Mb.

### Fraxinus spp

SNPs in candidate genes for *F. excelsior* were available from previous studies (Sollars et al. 2017; Stocks et al. 2019; Kelly et al. 2020). Of these, 2,500 associated with ash dieback disease susceptibility, 64 associated with emerald ash borer susceptibility, and 26 located in candidate regions linked to the *Oleaceae* family self-incompatibility system were selected. Filtering to discard triallelic SNPs and multiple hits on the BATG.0.5 ash reference genome resulted in 2,474 relevant SNPs (Sollars et al. 2017; https://www.ebi.ac.uk/ena/browser/view/GCA_900149125.1). Additionally, 324,590 SNPs were detected by mapping available transcriptome sequence data (Harper et al. 2016) against the BATG.0.5 genome, and filtering them for CG and AT variants, triallelic loci and multiple hits. The quality of a total of 327,064 SNPs was checked by TF, resulting in 85.4% (279,313) of them being classified as good quality (recommended).

Of these, a selection of SNPs was made for the screening array, with three priority levels: Priority 1 - SNPs (2,474) located in candidate genes (up to 8 SNPs per gene/transcript) associated with ash dieback disease resistance loci, and emerald ash borer susceptibility and incompatibility; Priority 2 - SNPs (62,713) from transcriptomic data, maximum of 8 SNPs per gene with a MAF > 5% ; Priority 3 - SNPs (92,253) from all other contigs in the genome sequence (Sollars et al., 2017) that were not included in Priorities 1 and 2. A total of 138,038 probes targeting 138,002 SNPs spanned 48,742 contigs, representing 753.8 Mb (86.9%) of the total genome assembly was included in the screening array (Sollars et al. 2017) (Figure 1, and Table S2). Recently, a contiguous reference genome BAT-G1.0 was released (Wood et al. 2025). The new assembly was used to locate the position of the SNPs. Flanking sequences of the probes were aligned with the procedure used for *Populus* (Figure 3).

**Figure 3:**
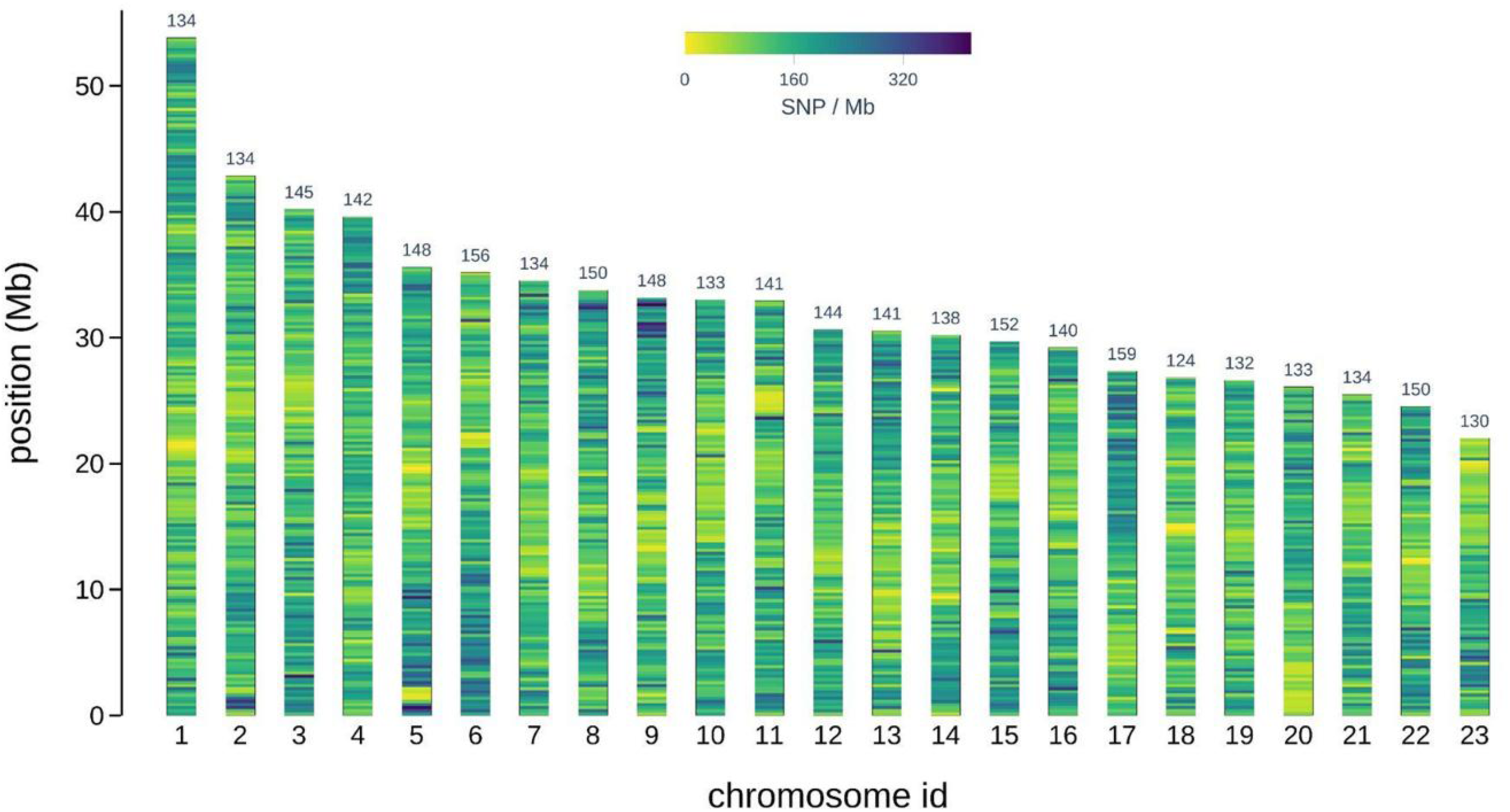
Fraxinus SNP density from the screening array on the BATG1.0 genome assembly of *Fraxinus excelsior* (Wood et al. 2025). The y axis represents the physical distance along each chromosome, split in 250 kb windows. The different colors depict SNP density. Over 76 % of the probes were successfully mapped along the *F. excelsior* BATG-1.0 genome assembly with the BLASTn tool using the following parameters: alignment coverage >0,9 and identity >0,95. The numbers above the chromosomes are the average number of SNPs per Mb.

### Pinus pinea

*P. pinea* is an orphan species and SNPs discovery was initiated during the B4EST project. Five individuals with distinct origins were selected from Spain, Greece, Lebanon, Turkey and Morocco in order to identify SNPs through RNA-Seq. RNA extractions and library preparations were performed according to standard protocols and carried out at IGATech (Udine, Italy). Sequencing was done on a HiSeq 2500 system (Illumina, San Diego, CA) in 125 bp paired-end mode. The read number ranged from 73 to 99 million/sample. After trimming and filtering, high-quality reads were aligned to the *Pinus taeda* v2.0 reference genome (Zimin et al. 2017) using the RNA-Seq STAR v2.6 aligner (Dobin et al. 2013) by setting the outFilterMultimapNmax parameter to 5 and the outFilterMismatchNmax to 6. Only reads with a mapping quality of at least 10 were selected using the samtools package (Li 2011) and duplicated sequences were removed using the picard tools MarkDuplicates utility (https://broadinstitute.github.io/picard/). Aligned reads were split into exon segments and sequences overhanging intronic regions were hard-clipped using the SplitNCigarReads tool of GATK v4.0.3.0 (DePristo et al. 2011). SNPs calling was performed using the HaplotypeCaller tool of GATK v4.0.3.0 (DePristo et al. 2011) with default parameters. SNPs were selected using GATK SelectVariants and filtered using GATK VariantFiltration by applying the following filter expression: QD < 2.0 || FS > 30.0. For each sample, SNPs were further filtered according to a minimum coverage of 8 and a minimum percentage of reads supporting the alternative allele of 20%. Finally, interspecific SNPs (i.e., *P. pinea* vs. *P. taeda* polymorphisms) were discarded and only positions informative in at least two samples were retained. A total of 128,279 SNPs with 35 bp upstream and downstream flanking sequences were obtained and submitted to TF for quality scoring. Most of them (121,572) were not recommended (94.7%). All recommended and neutral SNPs, along with approximately 30,000 additional non-recommended markers, were retained for the screening array and classified into four priority levels: Priority 1 consisted of 11,861 SNPs, including all recommended or neutral SNPs and additional isolated SNPs with flanking sequences with good sequence homology with *P. taeda* genome sequence (homology_count < 30); Priority 2 contained 11,957 SNPs with a probability of conversion > 0; Priority 3 included 7,395 SNPs that were targeted by a single probe, did not have interfering polymorphisms in their flanking sequences, and showed high homology with *P. taeda* genome sequence (wobble_count=0, homology_count<30 and probe_count=“empty”); Priority 4 consisted of 5,574 SNPs targeted by a single probe and without any interfering polymorphisms in the flanking sequences (wobble_count=0 and probe_count=“empty”) but not fulfilling the homology criteria required for Priority 3. Ultimately, 36,784 corresponding to 28,7% of the submitted SNPs were kept in the screening array design, including 5,871 A/T or C/G SNPs (Figure 1 and Table S2).

### Pinus pinaster

Most of the available SNPs came from genomic resources created in the framework of the GenTree project (https://www.gentree-h2020.eu; Milesi et al, 2024). They were discovered from an exome capture experiment carried out with a diversity panel of 163 individuals, with the same procedure as that described for *P. pinea*. Moreover, 5,165 high quality SNPs from the 9K Illumina Infinium array generated in Plomion et al. (2016) were added. Overall, 191,785 SNPs were sent for quality scoring. Most of them (92%) returned labelled not recommended. Five priority levels were defined to select SNPs : Priority 1 included 5,163 SNPs from the existing Illumina Infinium array; Priority 2 referred to 12,572 SNPs classified as recommended (3.4% of the markers) or neutral (4.1%); Priority 3 consisted of 31,106 SNPs with a probability of conversion > 0.3; Priority 4 consisted of 38,608 SNPs with no interfering polymorphisms and targeted by single probes (wobble_count=0 and probe_count=“empty”); Priority 5 consisted of 52,662 SNPs, with few mismatches in the probe (homology_count<=20). Among the selected 137,321 SNPs, 17,857 were A/T or C/G variants. Nearly all Priority 1 SNPs from the existing Infinium SNPs were tiled on forward and reverse strands to increase their chances of success (Figure 1 and Table S2). A total of 137,103 probes flanking 133, 408 SNPs were selected.

Ultimately, 449,688 SNPs for the three genus (*Pinus, Populus, Fraxinus*) were included in the screening array (Table S3).

### DNA extraction and SNPs genotyping

A panel of 476 trees (118 *Populus* spp., 131 *Fraxinus* spp., 108 *P. pinea*, and 119 *P. pinaster*), selected to represent the diversity in European breeding programs, were used for the screening array (Table S3 and S4). DNA was isolated from young plant material with different methods depending on the species (Table S4). DNA quantity and quality were estimated using a Nanodrop spectrophotometer and were sent to the TF genotyping lab (Thermo Fisher Scientific, Inc, Santa Clara, CA, USA) for genotyping. DNA concentrations were confirmed by Picogreen quantification (Table S4). Of the 476 samples only 14 had a DNA concentration below that recommended by TF, i.e. 10 ng/µl. These samples were included despite their lower concentration. Genotyping was conducted by TF according to standard protocols. SNP allele calling was done using the Axiom™ Analysis Suite (https://downloads.thermofisher.com/Axiom_Analysis_Suite_v_4.0.1_User_Guide.pdf). The genotypes were filtered using the Dish Quality Control metric (fluorescence signal contrast, computed on monomorphic loci) > 0.82 and according to the Quality Control Call Rate (as computed on high quality SNPs identified by TF on calibration plates) > 0.97. The software classified the SNPs into 6 categories: 1) PolyHighResolution (PHR), polymorphic SNPs leading to 3 well defined-genotype clusters; 2) MonoHighResolution (MHR), monomorphic SNPs leading to only one cluster of homozygous genotypes; 3) NoMinorHomozygous (NMH), one homozygous cluster is missing; 4) Call Rate Below threshold (CRBT); 5) Off-Target Variant (OTV); 6) Other: one or more QC metrics are below the thresholds.

## RESULTS

### Screening array performance

Genotype calling was firstly performed with the recommended default parameters for diploid species. Some of these default parameter values were subsequently adjusted for *Pinus* sp. and *Fraxinus* sp. In *Pinus* sp., call rate cut-off was lowered from 97% to 95%, as these species had fewer SNPs in the array, many of which had not been recommended by TF. In *Fraxinus* sp., the heterozygous strength offset (HetSO) parameter, which characterizes the position of the heterozygote cluster relative to the homozygote cluster, was lowered from -0.1 to -0.3.

We considered as “Informative SNPs” those included in the PHR category in which homozygotes and heterozygotes were both represented, and in the NMH category in which only one of the homozygote clusters was detected. MHR, i.e. monomorphic, was the third category. After filtering, 96% of the samples passed the quality control test: 100% of *P. pinea*, 93.3% of *P. pinaster*, 97.4% of *Populus* sp., and 96.2% of *Fraxinus* sp. The screening genotyping success rate ranged from 52.3% to 70.7% across species. For the two informative categories, i.e. PHR and NMH, lower success rates of 15.4% and 19.2% were obtained for *P. pinea* and *P. pinaster*, respectively. Much higher success rates were obtained for *Populus* (49.8%) and *Fraxinus* (47.4%) (Table 1).

**Table 1:** Performance of the screening array.

|  |  | *Pinus pinea* | *Pinus pinaster* | *Fraxinus spp* | *Populus spp* |
| --- | --- | --- | --- | --- | --- |
| Samples | Total number of samples | 108 | 119 | 135 | 118 |
|  | Passed samples | 108 | 111 | 126 | 115 |
|  | Av. cluster call rate | 99.65 | 99.48 | 99.51 | 99.58 |
|  | Sample Reproducibility |  |  | 99.94 | 99.64 |
| SNPs | Total nb of SNPs | 36,784 | 133,408 | 138,038 | 137,545 |
|  | PolyHighResolution (PHR) | 4,831 | 14,815 | 46,28 | 52,193 |
|  | NoMinorHom (NMH) | 840 | 10,853 | 19,081 | 16,266 |
|  | MonoHighResolution (MHR) | 20,329 | 50,26 | 27,651 | 3,407 |
|  | Call rate below threshold (CRBT) | 880 | 5,45 | 8,107 | 5,987 |
|  | Off target variant (OTV) | 518 | 3,755 | 7,557 | 18,998 |
|  | Other | 9,386 | 48,275 | 29,326 | 40,694 |
|  | % Genotyping success (PHR, NMH, MHR) | 70.68 | 56.91 | 67.38 | 52.25 |
|  | % Informative SNPs (PHR, NMH) | 15.42 | 19.24 | 47.35 | 49.77 |

### Genotype calling validation by comparison with existing genotyping data

For *Populus* species and *P. pinaster*, the availability of genotyping data obtained in previous studies for several genotypes allowed us to check the concordance between the platforms and assess the screening array performance (Faivre Rampant et al. 2016; Plomion et al. 2016).

### Concordance with available genotyping data and Mendelian segregation in Populus

We assessed the screening array performance by comparing previous DNA-Seq and RNA-Seq genotype calling data for 23 individuals that were also included in the screening panel (Table S5). The genotype concordance was between 91% and 97% for 20 genotypes. Two individuals, *P. nigra* 71077-308 and *P. deltoides* V5, exhibited lower concordance values of 87% and 80%, respectively. This could be explained by the low quality of their DNASeq and exome capture data. One last individual, Alaric, showed a very low concordance of 43,4%, suggesting that the genotyped sample represented a different, even if related, individual (Table S5).

We also compared the SNPs calling of 28 individuals genotyped with 6,737 SNPs shared between the Infinium array and the Axiom screening array. The genotype concordance exceeded 97% (Table S6). To test for Mendelian segregation, 5 trios (2 parents and one progeny) were included in the array. Moreover, 17 progenies whose parents had been genotyped via the array or previous genotyping tools were also present. Overall, we tested Mendelian segregation conformity in 25 trios, with the conformity expressed as the percentage of markers for which the descendant genotype could be correctly predicted from those of the parents. The conformities depended on the species of the parents, with *P. nigra* x *P. nigra* progenies showing the highest values (98%). *P. nigra* x *P. deltoides* hybrids, however, showed less conformity, with 6 to 14% non-concordant SNPs. This may have been partially due to the presence of null alleles for one of the parents that was not detected by the screening array. In that case, a heterozygous SNPs involving a null allele would be viewed as being homozygote. *P. deltoides* x *P. deltoides* progenies showed diverse conformity levels in the same range as that observed in *P. nigra* x *P. nigra* trios (Figure S1).

### Genotype calling validation in Pinus pinaster

We validated the genotyping data obtained with the screening array for a set of approximately 4,000 SNPs markers that were previously included in the Illumina Infinium Array (Plomion et al. 2016) on 70 *P. pinaster* trees genotyped by both technologies. The results showed a very high concordance rate for 59 of them (mean 98.9%), while the rates were much lower for the remaining 11 samples, with a mean of about 53%). The results obtained for these latter samples were probably due to identification errors, such as the accidental exchange of individuals or problems with the codes used to identify them (Figure S2).

### Selection of highly informative SNPs and the 4TREE array design Populus sp

Of the 137,545 targeted SNPs on the screening array, 52,193 PHRs were obtained and further selected in terms of their representativeness of the following categories: i) “Substitutions”, where each species is homozygote and hybrid heterozygote; ii) “Shared”, polymorphic in both *P. nigra* and *P. deltoides*; iii) “*P. deltoides*”, SNPs specific to *P. deltoides*; iv) SNPs specific to *P. nigra* and for which three subcategories were further defined: “*P. nigra*”, SNPs from the DNA-Seq and RNA-Seq data, “Illumina array”, SNPs from the Infinium black poplar array, and “GWAS”, SNPs from GWAS studies related to wood properties. After SNPs assignment to each category, those with inconsistent Mendelian segregation were removed. SNPs equally distributed along chromosomes were carefully selected in each species, while conserving SNPs of interest (Figure 4; Figure 5). For *P. nigra* priority was given to SNPs included in the Illumina array. Eight SNPs classified within the NMH as being associated with wood properties in GWAS studies were also included in the selection. An additional filter, based on density criteria was applied to reach a total of 15,753 SNPs evenly distributed on the genome.

**Figure 4:**
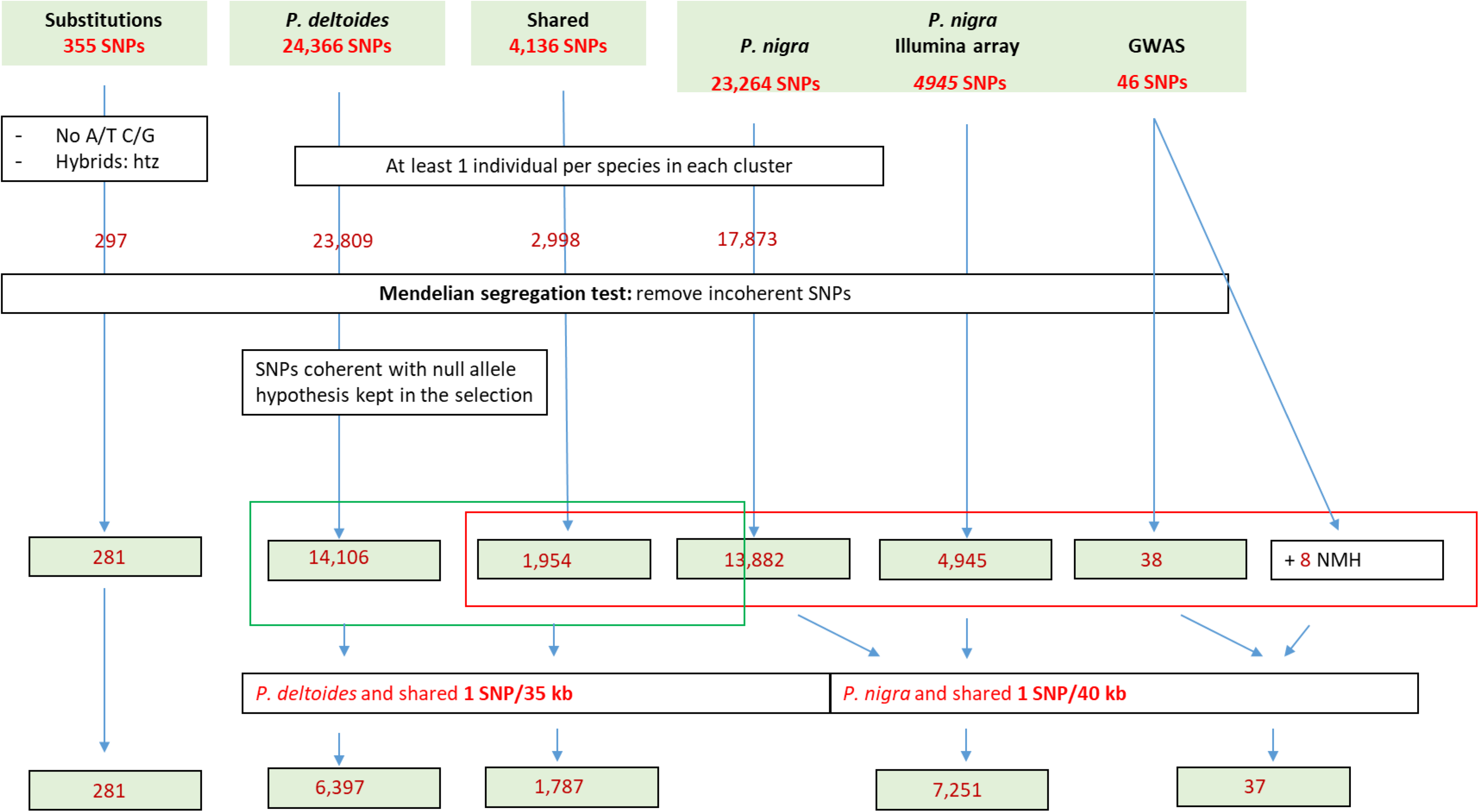
Diagram outlining the number of SNP at each SNPs selection step for the design of the *Populus* sp. 4TREE array part

**Figure 5:**
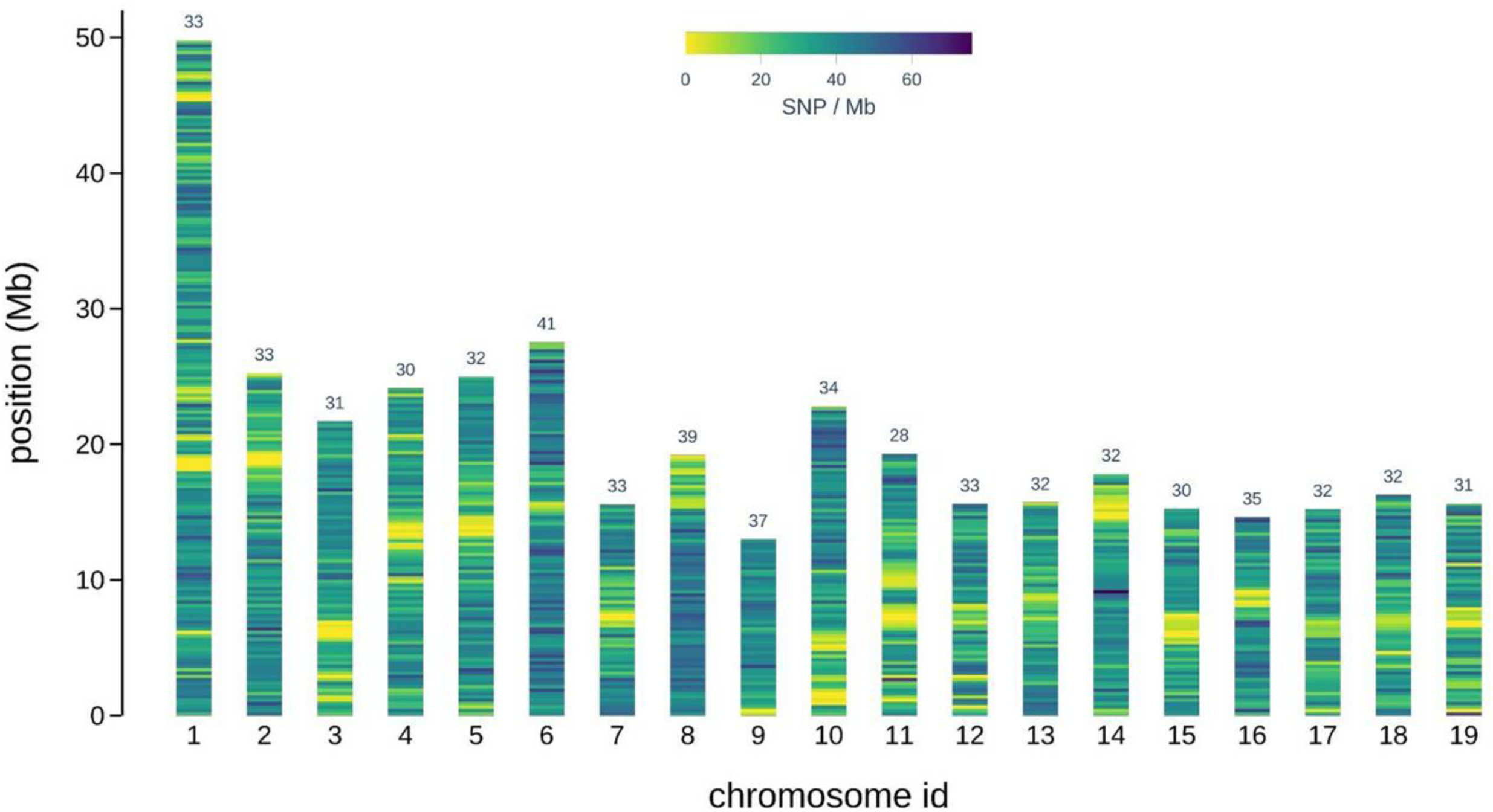
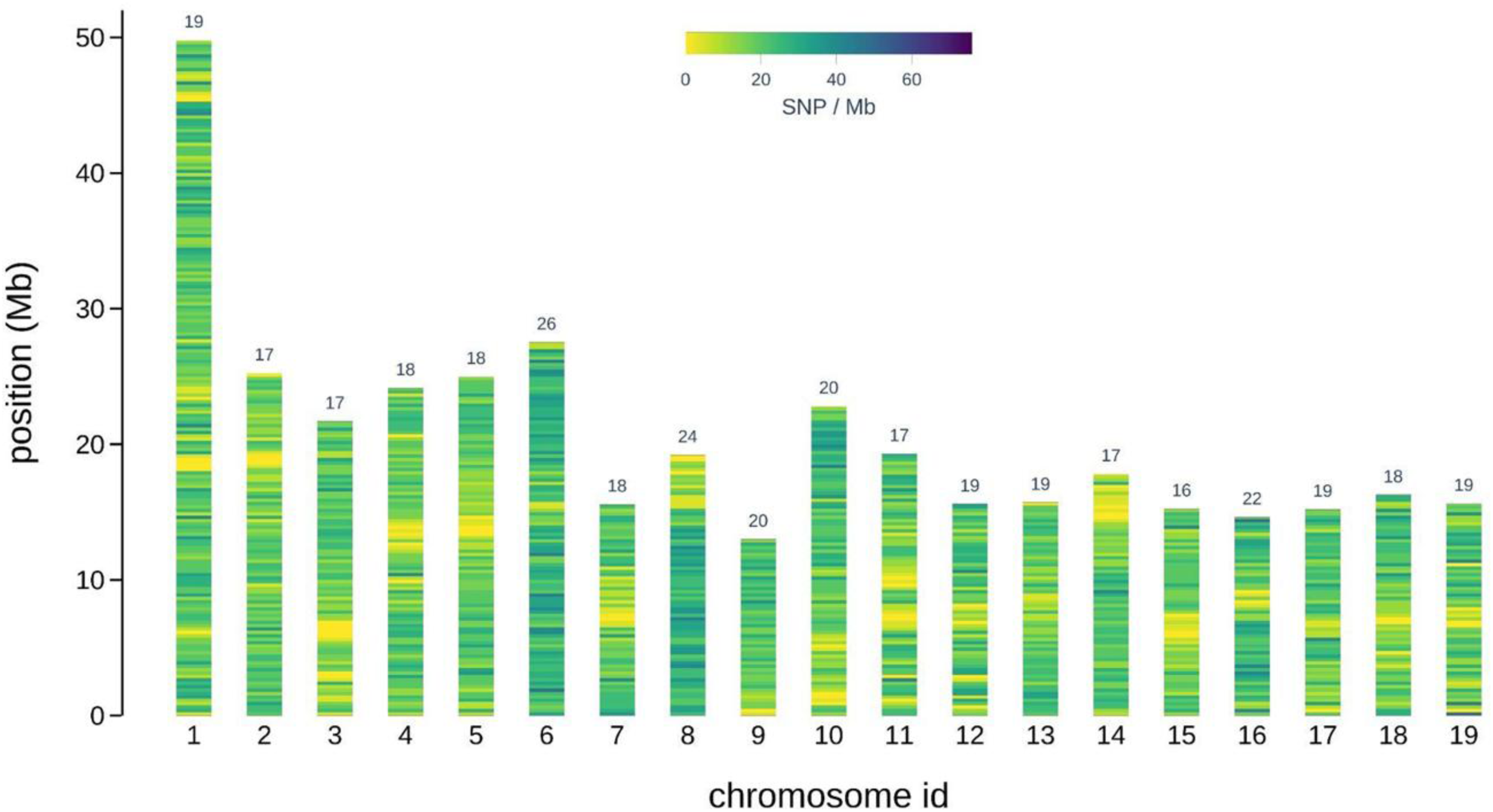
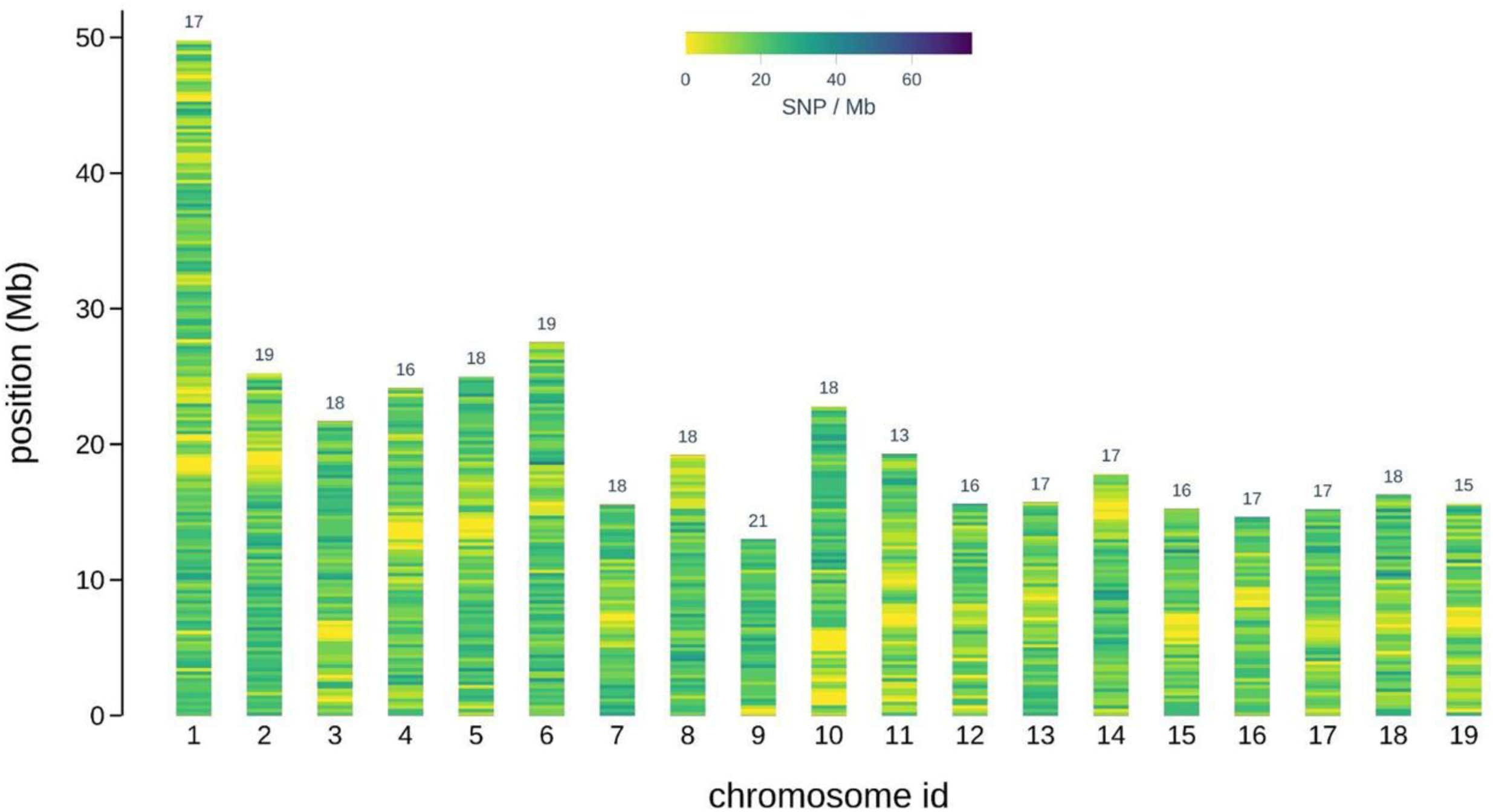
Distribution of the selected SNP of the 4TREE array for *Populus spp.* on the *Populus trichocarpa* genome V4.0. a) SNP density along chromosomes ; b) number of *P. nigra* SNP on each of the 19 chromosomes; c) number of *P. deltoides* SNP on each of the 19 chromosomes. The numbers above the chromosomes are the average number of SNP per Mb.

### Fraxinus spp

Priority 1 for the *Fraxinus* species was to integrate SNPs associated with traits of interest. These 1,704 SNPs were selected among the PHR, NMH, and MHR candidates (the rest of the markers submitted were not recommended by Thermo Fisher and not included); 51 SNPs were associated with emerald ash borer resistance/susceptibility, 1,633 with dieback disease, and 20 were from contigs supposedly associated with the *Oleaceae* self-incompatibility system. Priority 2 consisted of the 200 SNPs with highest loading for the first Principal Component of a PCA conducted on a set of accessions including 5 *F. angustifolia* and 111 *F. excelsior* (putative hybrids were screened out) as this first PC obviously separated the two species (Table S7). Priority 3 aimed at achieving even coverage of the *F. excelsior* draft genome using PHRs only, with 1 SNPs per contig. This last category consisted of 15,620 SNPs. A second SNPs per contig (Priority 4) was also included whenever possible. Overall, 17,524 SNPs were selected for *Fraxinus* spp. Re-mapping of these robust SNPs resulted in 23 chromosome hits, 2 mitochondrial scaffolds and 52 unplaced scaffolds.

### Pinus pinea

The majority of *P. pinea* SNPs (20,329) were classified as MHR as they were monomorphic in the analyzed dataset. An additional 10,784 markers did not pass the standard Axiom quality filtering step, while the remaining 5,671 were polymorphic, including 4,831 that were classified as PHR, while the remaining 840 were ranked in the NMH category. All markers classified as PHR and NMH (5,671 overall) were selected due to the scarcity of available polymorphic markers.

### Pinus pinaster

The majority of *P. pinaster* markers used for the screening (57,480) did not pass the standard Axiom quality filtering, while 50,260 additional markers were classified as MHR. Among the remaining SNPs, 25,668 were polymorphic and 14,815 of these were PHR and 10,853 were NMH. The 16,003 *P. pinaster* markers selected for the 4TREE array were classified into three priority levels: i) Priority 1 – 3,889 PHRs (3,441) or NMHs (416) that had already been included in a previous Illumina Infinium array; ii) Priority 2 – 8,160 PHRs that were located at least 100 bp apart; and iii) Priority 3 – 3,954 NMHs located at least 100 bp away from other selected markers.

### 4TREE array composition

Finally, a total of 54,952 SNPs were sent to TF for a last round of scoring and optimization in order to find the best combination for the 4TREE array design. The final array consisted of 45,893 SNPs, including 13,408 *Populus* sp. SNPs (280 substitutions, 932 shared, 5,734 *P. deltoides*, 6,462 *P. nigra* (2407 Illumina, 34 GWAS); 13,407 *Fraxinus* sp. SNPs; 5,671 *P. pinea* SNPs and 13,407 *P. pinaster* SNPs.

This final array was the result of an optimized trade-off between the four species and the quality of available markers. The list of SNPs and their flanking sequences can be found in table S8. The majority of the 13,408 Populus’s SNPs (97%) was located on the genome *P. trichocarpa* V4.0. The density of *P. nigra* SNPs per Mb ranged from 16 on chromosome 18 to 25 on chromosome 8. Density of *P. deltoides* SNPs varied from 14 SNPs per Mb on chromosome 19 up to 23 on chromosome 9 (Figure 5). Among the 13,407 F*raxinu*s SNPs, 91.5% were uniquely mapped on the newly assembled chromosomes, with the number of SNPs per chromosome ranging from 801 (for the longest chromosome) to 231 (for the shortest chromosome). On average, we obtained an SNPs density of 15.72 SNPs per Mb, with a minimum density of 12.6 SNPs per Mb (chr 18) and a maximum density of 21.65 SNPs per Mb (chr 23) (Figure 6, Table S8)

**Figure 6:**
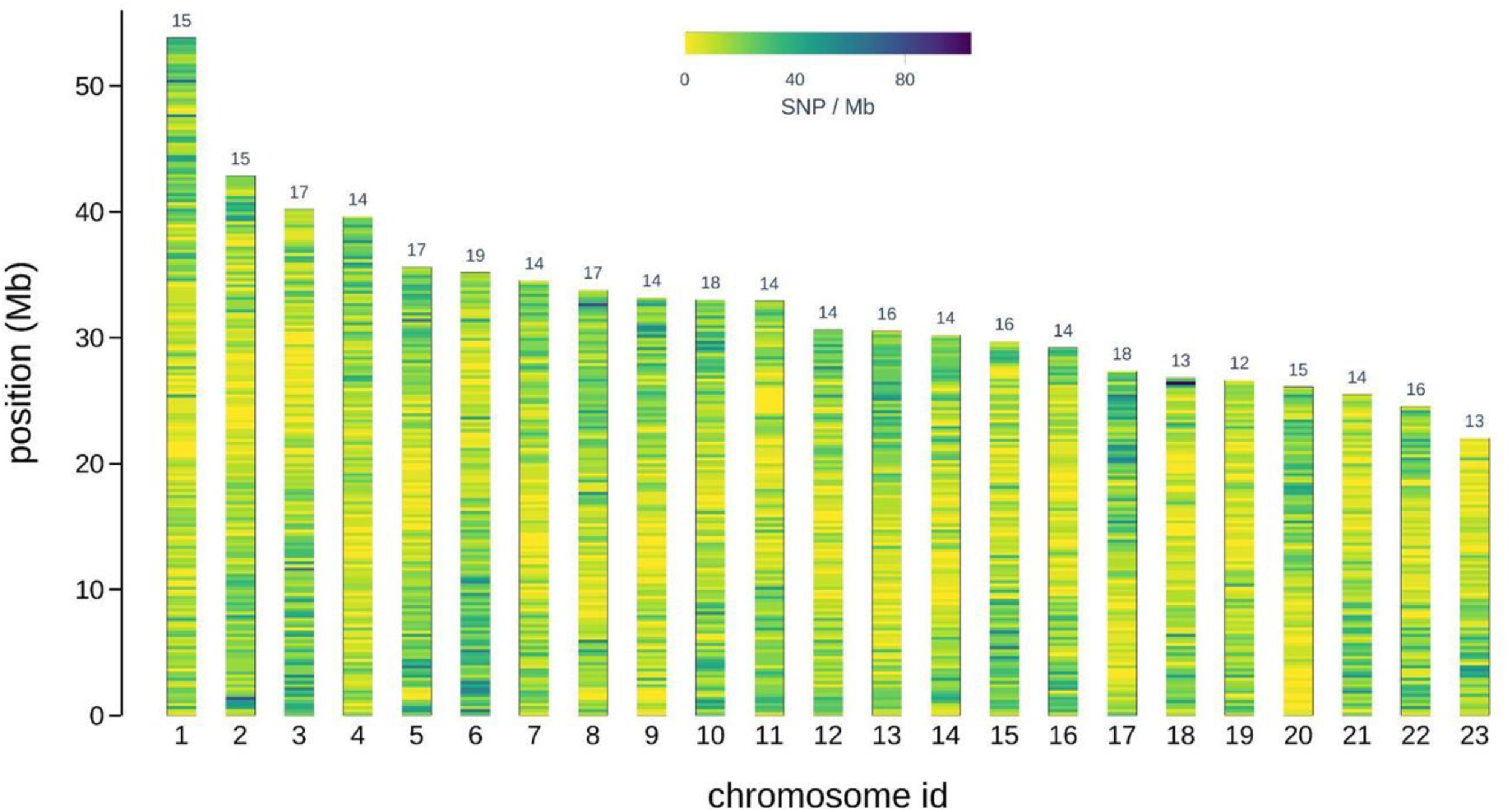
Distribution of the selected SNPs of the 4TREE array for *Fraxinus* spp. on the BATG1.0 genome assembly of *F*. *excelsior*. The numbers above the chromosomes are the average number of SNP per Mb.

### 4TREE array validation

To evaluate the performance of the 4TREE array, genotyping results of 2,229 *Populus*, 2,088 *Fraxinus*, 1,897 *P. pinea*, and 1,858 *P. pinaster* individuals were obtained according to the screening array test procedure (Table 2).

**Table 2:**
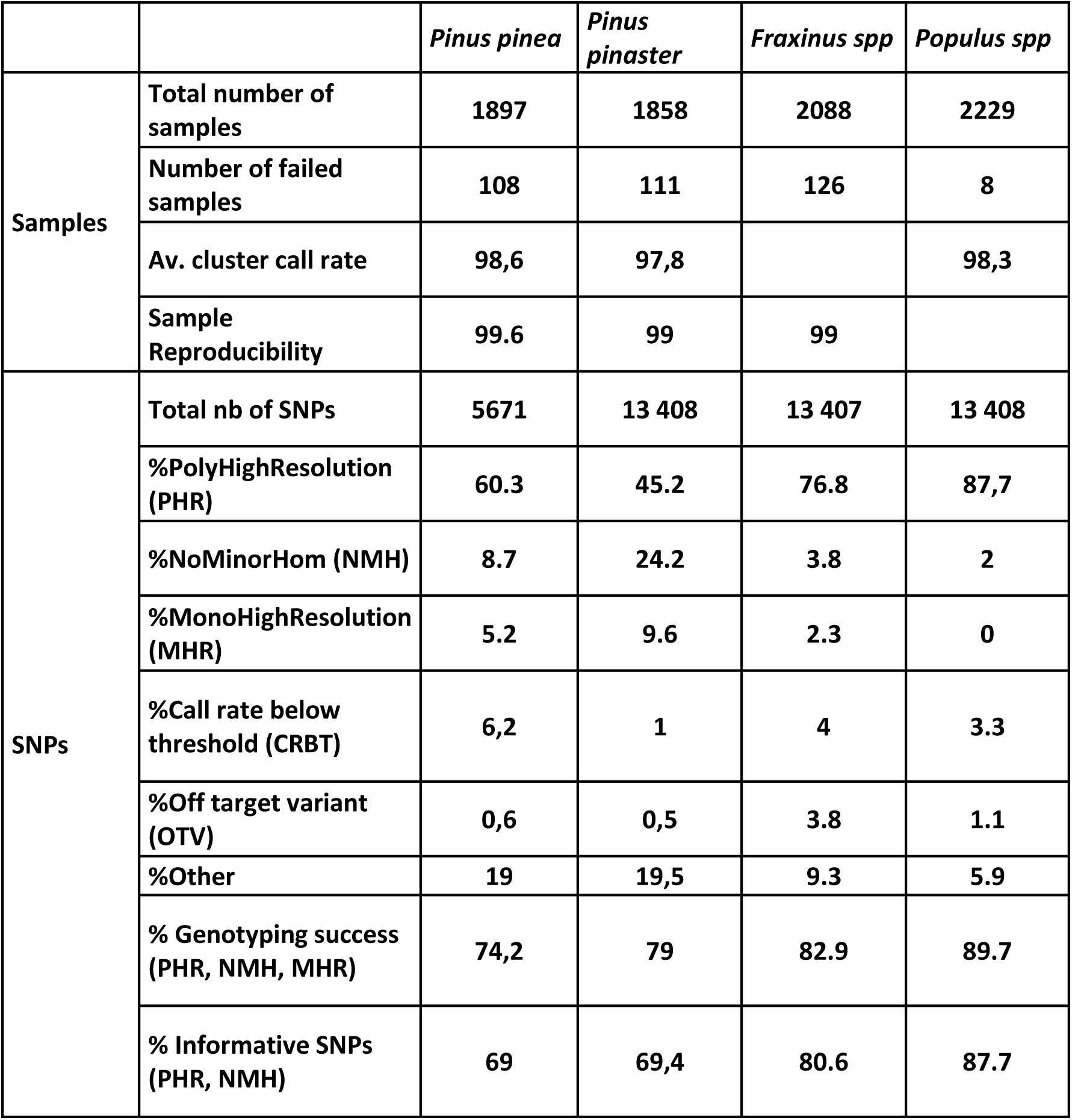
Performance of the 4TREE array SUPPLEMENTARY TABLES.

The 2,229 poplar trees included 1,602 *P. nigra*, 174 *P. deltoides*, 79 *P. x canadensis*, and 66 *Populus* spp. Only 8 samples could not be genotyped as they did not pass the TF quality control threshold > 0.82. Concerning the genotyping results, SNPs were classified into the 6 classes proposed in the Axiom best practices guide. The genotyping success rate was 89.7%. Only 2% of the markers were not informative and belonged to the MHR class, while the remaining 10.3% included the useless classes (other 5.9%, CRBT 3.3%, OTV 1.1%). To assess the repeatability between the two arrays (screening and final), the concordance between 5 poplar genotypes included in the two arrays was measured. This analysis revealed that the concordance of the data for all samples analyzed by both arrays was 100% (Figure S3). Similarly, the analysis of 68 technical replicates of *P. deltoides*, *P. nigra*, and *P.* x *canadensis* lines showed that the concordance of the genotypes included in the 4TREE array ranged from 98.9% to 100%, thus corroborating the high performance of the multi-species array developed in the present study (Figure S3).

A majority of the 2,088 *Fraxinus* spp. samples were represented by *F. excelsior* individuals. About 100 *F. angustifolia* samples and some potential hybrids between the two species were also present. Overall, 76.8% of the 4TREE array SNPs were classified as PHR, while 3.8% were NMH. The remaining 19.4% (call rate < 4% threshold, MHR 2.3%, OTV 3.8%, and Other 9.3%) was not informative for the panel under study. A very high genotype call concordance rate (> 99%) was found for 8 pairs of replicated samples. All samples previously classified as *F. angustifolia* based on morphological traits were correctly assigned by the array using a principal coordinate analysis (PCoA). In addition, of all of the samples labelled as *F. excelsior*, 19 were positioned between the *F. excelsior* and *F. angustifolia* clusters in the PCoA, indicating they were potential hybrids between the two species (data not shown). For both *Pinus* species, most of the analysed samples were successfully genotyped (98.8% and 98.2% for *P. pinea* and *P. pinaster*, respectively). For the two species, the proportion of successfully genotyped markers was similar, with 74.3% for stone pine and 79% for maritime pine, although for stone pine a higher fraction of PHRs was found (60.3%). Consequently, a lower fraction of NMHs (8.7%) and MHRs (5.2%), in comparison to maritime pine (45.2% PHR, 24.2% NMH, and 9.6% MHR), were obtained. We assessed the genotyping reproducibility by genotyping twice five *P. pinea* samples and nine *P. pinaster* samples. For both species, we obtained a high concordance between the genotypes called in the two replicates, with an average of 99.6% in stone pine and 99% in maritime pine.

## DISCUSSION

The aim of this European collaborative work was to create a genotyping tool to improve genomic research on the genetic resources of four important forest species. The main objective was to develop a dependable, robust and reusable tool that would enable data from various studies to be compared and combined, even after the B4EST project had ended. To this end, we opted for the SNP array platform, well known for its robustness and reproducibility with low requirements of bioinformatic expertise. The operating costs of these tools depend on the number of SNPs and individuals under study. The higher these numbers, the lower the cost per data point. Unlike field crops, it is difficult to obtain a combination of SNPs and individuals large enough to be cost-effective for a single-species array for most forest species (Rasheed et al. 2017). However, there are exceptions among the most commercially traded forest species, such as *Eucalyptus* sp., *Pseudotsuga menziesii* (Howe et al. 2020), *P. sylvestris* (Kastally et al. 2022), *P. abies* (Bernhardson et al. 2021), *P. radiata* (Graham et al. 2022), and *P. taeda* (Caballero et al. 2021). For most forest tree species, the number of SNPs may not be high because of the lack of genomic resources, e.g. in our study for *P. pinea*, and/or because of the low number of individuals available for testing. Reduced genotyping costs for species with limited genomic resources or demand can be achieved by creating a common SNP array for several species (Montanari et al., 2022). Our study exemplifies this approach, in line with findings on other forest species (Silva-Junior et al. 2015; Perry et al. 2020; Jackson et al. 2021). We have also demonstrated the effectiveness and relevance of exploiting synteny between closely related species in the development of genotyping tools. Furthermore, we have shown that the 4TREE matrix is useful for genotyping different species belonging to the *Populus* and *Fraxinus* genera.

The conversion rate (PHR, MHR, and NMH) obtained with the screening array ranged from 52% to 71%. Unexpectedly, the lowest rate was obtained in poplar (52%) and the highest rate in *P. pinea*. This reduced poplar value conversion rate could be explained by several factors depending on the species and quality of reference genome assembly. For poplars, SNPs for *P. deltoides* and *P. nigra* were selected using the *P. trichocarpa* reference genome (v4.0), which may have affected SNP conversion rates, as sequence divergence between reference and target species can impact probe hybridization efficiency. Otherwise for ash SNP discovery relied on a highly fragmented reference genome, which may have limited accurate SNP positioning and flanking sequence selection, thereby reducing probe performance and SNP conversion rates. These results highlighted the importance of having a high quality assembled reference genome available, not only for SNPs identification but also for the probe design efficiency. For *P. pinaster* and *P. pinea*, two conifer species originating from Southern Europe, the selection of SNPs and flanking sequences was performed using, as reference, the genome of *Pinus taeda*, a North American species, the most complete conifer genome, considered as a genome reference for conifers, publicly available at the beginning of the project. A high polymorphism between the genomes of these conifer species could be hypothetized. The conversion rate obtained in *P. pinaster* (56%) is lower than that observed in *P. sylvestris* (Kastally et al. 2022; Estravis-Barcala et al. 2024) and *Picea abies* (Bernhardsson et al, 2021), but higher than that obtained in *P. radiata* (Graham et al, 2022). The high MHR rate in the two *Pinus* species could be due to the location of the selected SNPs within genes, as exemplified when comparing a coding-regions derived and a genome-wide derived SNPs Axiom arrays in *P. sylvestris* (Kastally et al. 2022; Estravis-Barcala 2024). The same trend has also been observed in *P. radiata* (Graham et al. 2022)

We assessed the performance of the SNPs on the screening array by testing the concordance with other genotyping data in poplar and maritime pine. In the case of poplar, the concordance with transcriptome, resequencing and microarray data showed a rate equivalent to the one recorded in *P. nigra* (Faivre-Rampant et al. 2016; Rogier et al. 2018), and in other species such as strawberry (Bassil et al. 2015), apple (Bianco et al. 2016), and cacao (Livingstone et al. 2017). The concordance rate between the maritime pine genotypes in our study and those obtained by Plomion et al. (2016) was > 98%, which was the same rate as obtained in apple (Bianco et al. 2016), poplar (this study), and eucalyptus. In poplar, we were also able to test for Mendelian segregation. The concordance obtained in *P. nigra* intraspecific crosses was 98%, equivalent to that observed in a previous study (Faivre Rampant et al. 2016) and in *Picea abies* (Bernhardsson et al. 2020).

The two-steps strategy proposed by TF has turned out to be a winning approach for the selection of the most informative SNPs for advanced genetic studies. The conversion rate we obtained with the 4TREE array was 89.7% for *Populus* spp. and 80% for *Fraxinus* spp., while the rates obtained in *Pinus* species were 74.3% for *P. pinea* and 79% for *P. pinaster*. The approach has also been successfully applied in pear (Montanari et al. 2019), *P. abies* (Bernhadsson et al. 2020), *P. taeda* (Caballero et al. 2021), *P. sylvestris* (Kastelly et al. 2021), *P. radiata* (Graham et al. 2022), and tropical pines (Jackson et al. 2022).

The array is already being used by the B4EST project partners for studies on diversity in genetic resources (natural and breeding populations), genetic mapping and association genetics (Archambeau et al. 2023). The new 4TREE array enables more precise clonal identification and identity verification (post-hoc traceability) than the SSR markers currently used in *P. pinea* (Olsson et al. 2025). Moreover, it enables the detection of population structure with greater resolution in the westernmost part of the *P. pinea* distribution range and allows for accurate kinship assignment and pedigree reconstruction (Olsson et al. 2025). Similarly, the 4TREE array has been found useful in detecting a more refined distribution of the genetic diversity of *P. pinaster* across its distribution range and identifying new gene pools that were not previously detected using other molecular markers. This information has been used to improve our understanding of the biogeographic history (Theraroz et al. 2024).

Genomic promises to speed up breeding schemes, while reducing costs (Calleja-Rodriguez et al. 2020; Grattapaglia 2022). In this context, this array has enabled the construction of new, high-performance predictive models for genomic selection in *P. pinaster* (Papin et al 2024a; 2024b).

Furthermore, the 4TREE array was used outside the B4EST consortium to identify SNPs associated with pine wilt disease response in *P. pinaster* (Miguel, pers. Com.). The chip is also being used by poplar foresters to identify wooly poplar aphid resistance loci using two linkage mapping approaches based on two different F_1_ *P. deltoides* x *P. nigra* and *P. nigra* x *P. nigra* mapping populations (Biselli et al. in preparation). Recently, in the frame of FORGENIUS Consortium, Pinosio et al. (2025) showed that 3,031 *Populus* SNPs were useful to genotype 126 *P. alba* trees. The authors demonstrated that the 4TREE array could be used as a genotyping tool to characterize European *P. nigra* and *P. alba* populations. A recent implementation of the 4TREE array (Szukala et al. 2025) failed at detecting SNPs associated with ash tolerance to ADB possibly due to polygenic architecture of the trait combined with rapid decay of LD. It however succeeded at detecting interspecific hybrids and at revealing a genetic structure that follows a geographical pattern.

Nevertheless, the use of SNP arrays in population genomics involves a number of trade-offs that need to be discussed. First, SNP arrays are inherently biased toward previously identified polymorphisms, leading to an ascertainment bias that can distort allele frequency spectra, reduce estimates of rare variants, and affect inferences of demographic history and population differentiation. This bias is particularly pronounced when arrays are developed using a limited number of discovery populations or reference species, as is often the case in forest trees. Second, compared with Genotyping by sequencing, the fixed nature of SNP arrays prevents the detection of novel or population-specific variants, structural variants, and copy number variations, which may play an important role in local adaptation (Scaglione et al. 2019). Third, reduced marker density per species in multispecies arrays can limit genome coverage and mapping resolution, potentially decreasing power for genome-wide association studies (GWAS) or genomic selection in species with large and complex genomes. These limitations highlight the importance of carefully matching the choice of genotyping technology to the biological questions addressed, while emphasizing that SNP arrays remain a powerful and cost-effective tool for large-scale, standardized population genomic analyses (Werghi et al, 2025).

The 4TREE array was the result of a highly collaborative effort between multiple partners to compile the best resources available regarding the *P. pinaster*, *P. pinea*, *Populus* sp, and *Fraxinus* sp in a cost-effective genotyping tool. Although the multiple species array concept automatically implies a reduced number of markers per species, the fact of going through a preliminary SNPs screening phase generated higher quality results in the final allotment per species. Moreover, the real benefit of a shared tool must be found undeniably in the larger panel of potential users, which in turn can make the product competitive in terms of costs and extend its availability over time. This strategy could be applied to other species that are of scientific interest but have initially low demand, providing cost-effective genotyping access for many so-called orphan species. The latter prospect could be encouraged as part of efforts to diversify the species of interest in order to address the growing challenge of climate change. Last but not least in terms of benefits, mutualizing expertise across species and partners is especially relevant given that bioinformatics expertise, for example, can be scarce or even unavailable in some institutions for some species.

## ACKNOWLEDGEMENTS

We thank F. Isik from North Carolina State University, Raleigh, NC (USA) for leading the Conifer Scientific Consortium and for negotiations with ThermoFisher. We would also like to thank M. L. Schneider from ThermoFisher for facilitating all the exchanges between B4EST partners and the USA-Santa Clara ThermoFisher Microarray Research Services Lab. The authors acknowledge Genoscope (CEA-IbFJ) for providing computational resources.

## AUTHOR CONTRIBUTIONS

Conceptualization: J. Buiteveld, P. Copini, M.J.M. Smulders, G. Nervo, L. Cattivelli, A. Dowkiw, G.G. Vendramin, V. Jorge, L. Sanchez, S. C. González-Martínez, C. Bastien, P. Faivre Rampant

Resources: G. Nervo, L. Vietto, G.G. Vendramin (species leader for *P. pinea*), A.Dowkiw (plant material selection and field collecting, species leader for Ash), V. Benoit, C. Buret (field collecting, DNA extraction), F. Bagnoli, I. Spanu (DNA extractions), S. C. GonzálezMartínez (field collecting and species leader for *P. pinea* and *P. pinaster*), R.J.A Buggs, L. J. Kelly, C.L. Metheringham (candidate SNP loci and genomic data for *Fraxinus*), V. Jorge, V. Segura, S. Pinosio, P. Faivre Rampant.

Methodology : P. Copini, J. Buiteveld, D. Esselink, L. Kodde, G. Tumino, M.J.M. Smulders, C. Biselli, A. Fricano, G.G. Vendramin, O. Rogier, R. Guibaud, S. Pinosio, P. Faivre Rampant.

Formal analysis: J. Drouaud, D. Esselink, L. Kodde, G. Tumino, A. Fricano, S. Ben-Sadoun, O. Rogier, R. Guibaud, S. Pinosio.

Writing original draft: R. Guilbaud, Patricia Faivre Rampant, M.J.M. Smulders, G. Tumino, G.G. Vendramin.

Writing review editing: R. Guilbaud, P. Faivre Rampant, J. Buiteveld, J. Drouaud, G. Tumino, C. Biselli, A. Fricano, L. Cattivelli, G. Nervo, A. Dowkiw, G.G. Vendramin, L. Sanchez.

## SUPPLEMENTARY FIGURES

Figure S1: Mendelian inheritance of Genotyping results in *Populus* trios

Figure S2: Concordance rate obtained between the 4TREE array genotyping results and those obtained with Illumina for *Pinus pinaster* (Plomion et al. 2016).

Figure S3: a) Concordance between 5 poplar genotypes included in the screening and the 4TREE arrays. b) Repeatability between technical replicates for 68 poplar individuals in the 4TREE array. D, *P. deltoides*, N, *P. nigra*, DN, *P. canadensis*

Table S1: List of published forest tree SNPs arrays at the beginning the B4EST project Table S2: Affymetrix SNPs scoring features

Table S3: Number of SNPs per species in the screening array and number of individuals used to assess the screening array

Table S4: Plant material used to assess the screening array and associated metadata

Table S5: Concordance rate obtained between the screening array results and DNA/RNA-seq data in *Populus* sp.

Table S6: Concordance rate obtained between the screening array results and the Infinium Black Poplar array genotypes (Faivre Rampant et al. 2016)

Table S7: List of *Fraxinus angustifolia* SNPs probes

Table S8: List of 4TREE array SNPs and their flanking sequences

