## Supplementary figures and images for "Development of the 4TREE SNP array, a forest multispecies array to enhance European Breeding and conservation programs in pine, poplar and ash"

### Supplemental Figures 1-3

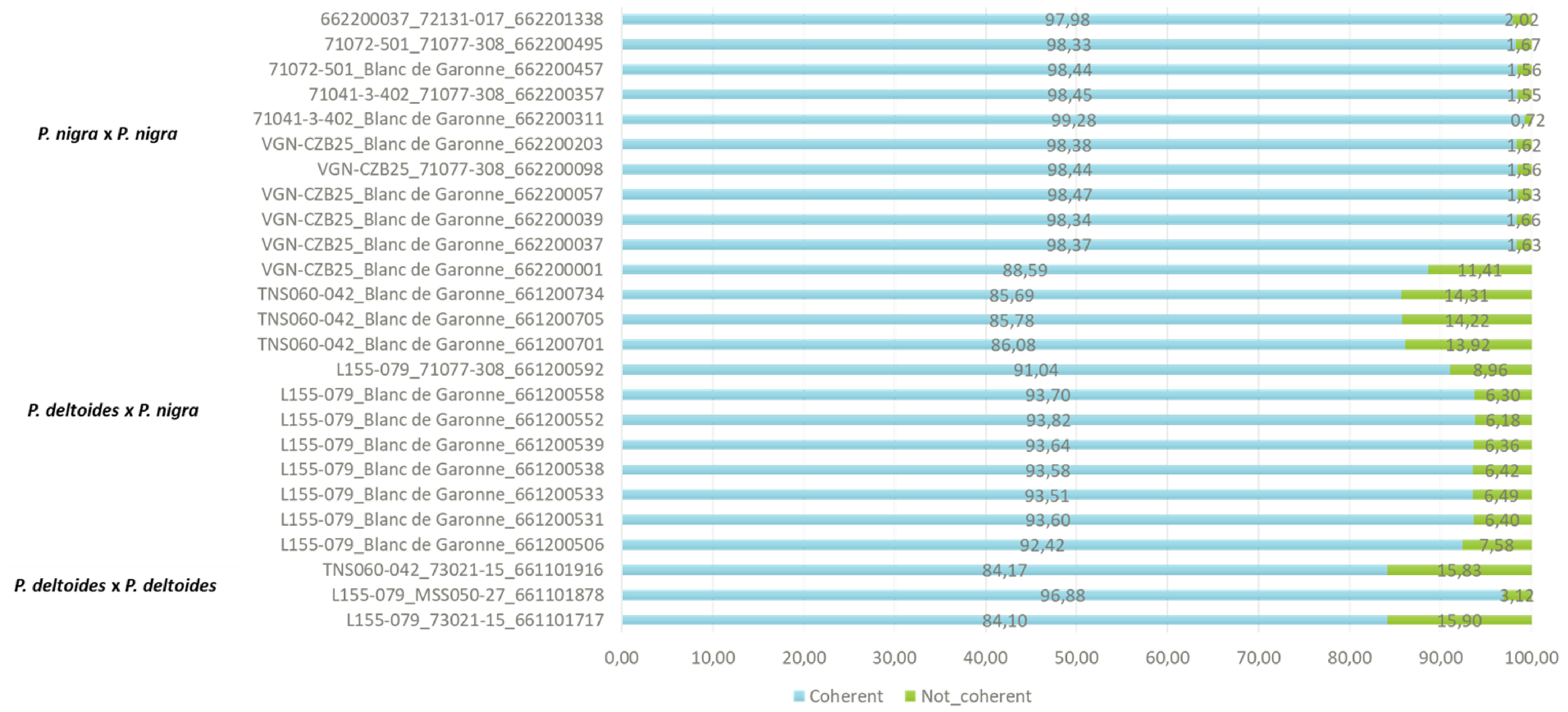

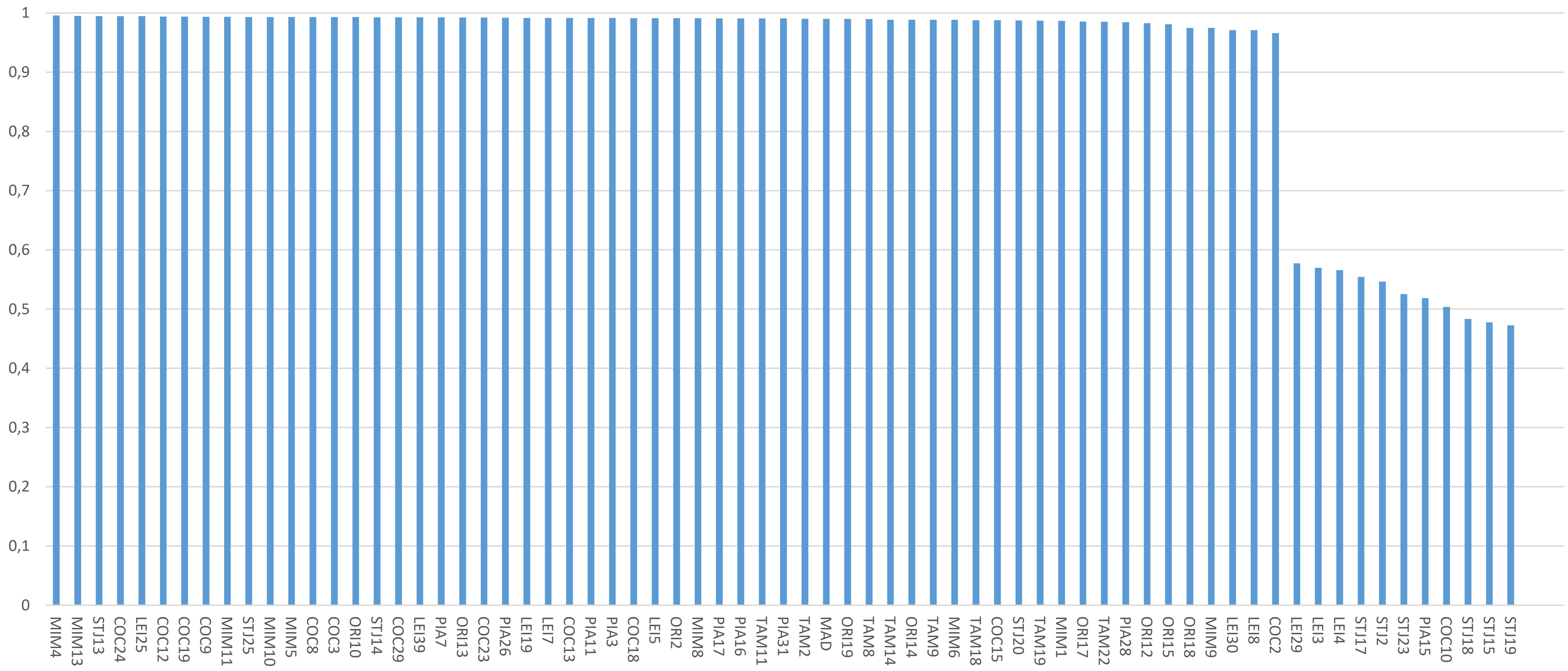

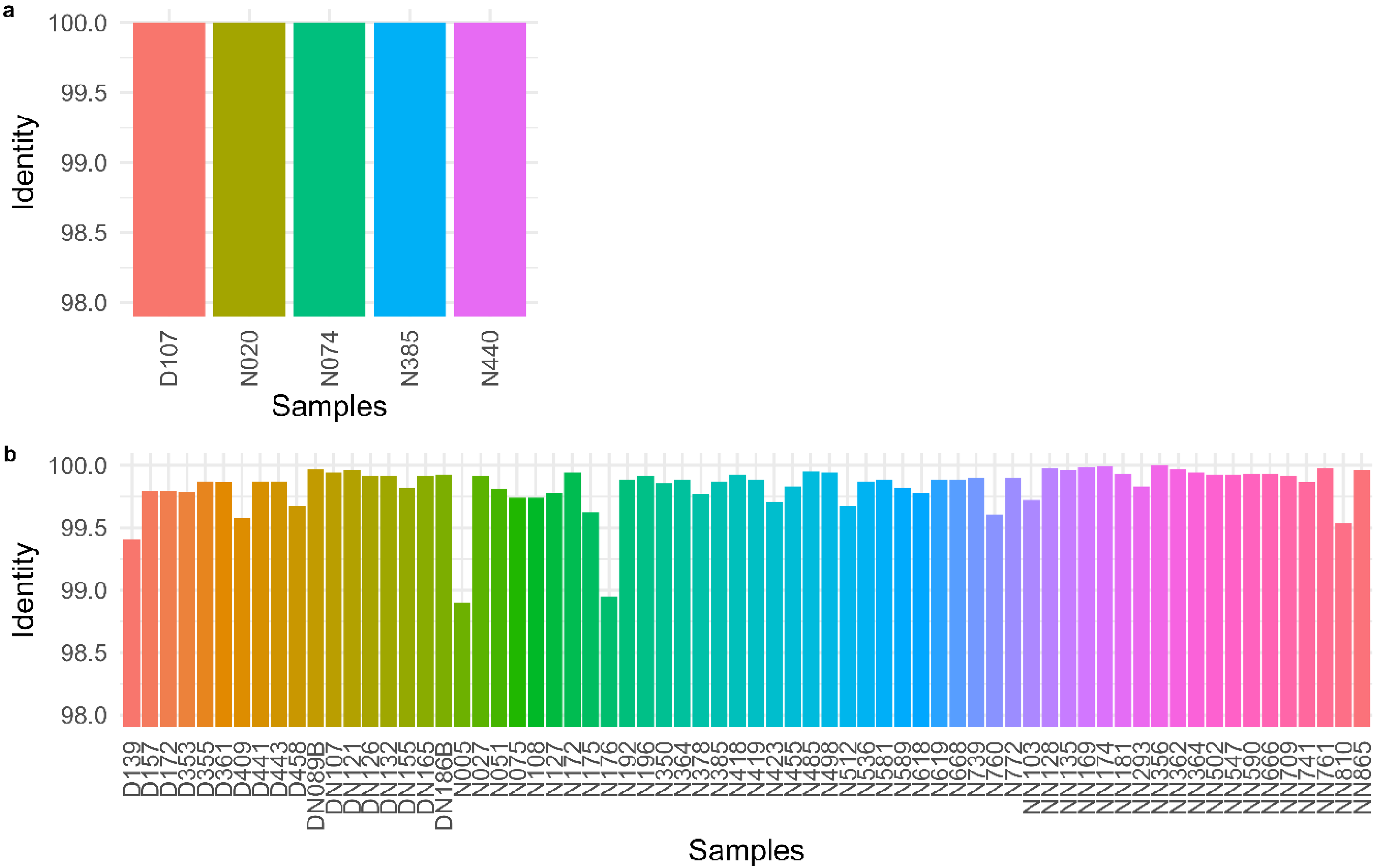
